# Identifying Strong Modulators of Cellular Quiescence Depth Across Different Quiescent Cells and Conditions

**DOI:** 10.1101/2022.11.19.517178

**Authors:** Eric Lu, Guang Yao

## Abstract

The proper balance and transition between cellular quiescence and proliferation are critical to tissue homeostasis, repair, and regeneration. The likelihood of quiescence-to-proliferation transition is inversely correlated with quiescence depth, and deep quiescent cells are less likely to exit quiescence and reenter the cell cycle than shallow quiescent cells. The regulatory mechanisms of quiescence depth are poorly understood but essential for developing strategies against hypo- or hyper-proliferation diseases such as aging and cancer. Our earlier studies have demonstrated that the activation threshold of the bistable Rb-E2F gene network switch (Th_E2F_) controls quiescence depth. We have also identified coarse- and fine-tuning Th_E2F_ modulators in rat embryonic fibroblasts. To examine whether other quiescent cells (including most adult stem and progenitor cells) under different environmental conditions use the same or different modulators of quiescence depth, here we studied the behaviors of 30,000 theoretical quiescent cell models that each support a functional Rb-E2F bistable switch with a unique parameter set. We found that although the vastly heterogeneous quiescent cell models exhibited no apparent parameter patterns, they converged at two alternative groups of strong quiescence-depth modulators (G1 cyclin/cdk-related and Rb/E2F complex-related). Our further machine learning (decision tree) analysis suggested that the Rb protein level and dephosphorylation rate in quiescent cells determine which modulator group to use to regulate quiescence depth.

## INTRODUCTION

Quiescence is a cellular state where division is halted but reversible upon growth stimulation or the removal of inhibitory signals. Many cells in the human body, including most adult stem and progenitor cells, are quiescent. Quiescence provides a safeguard mechanism for long-lived cells against environmental stressors and damages accumulating during cell proliferation cycles^1,2^.

Quiescence is not a homogeneous state but rather heterogeneous with varying depths^3,4^. Deep quiescent cells require stronger stimulation and take longer to reenter the cell cycle than shallow quiescent cells. Quiescence depth increases with longer culture duration *in vitro*^5–11^ and with aging *in vivo*^12,13^, whereas it decreases as seen in stem cells primed for quiescence exit upon tissue damage or alerting signals^14–17^. Conceivably, quiescence depth determines the proliferative potential of quiescent cells; dysregulation of quiescence depth can lead to hyper- or hypo-proliferative diseases, such as fibrosis, autoimmune disease, cancer, and aging.

We have previously shown that the Rb-E2F pathway functions as a bistable switch to convert transient and graded serum growth signals into an all-or-none E2F activation and cell fate transition from quiescence to proliferation^4,18,19^. We also found that quiescence depth is regulated by the activation threshold of the Rb-E2F bistable switch (Th_E2F_)^20^. Th_E2F_ refers to the minimum serum concentration at which the Rb-E2F bistable switch in quiescent cells turns from the E2F-off state to E2F-on. A higher Th_E2F_ corresponds to deeper quiescence.

What regulates Th_E2F_ and thus quiescence depth? By simulating the effects of parameter perturbations and experimental tests, we have identified groups of coarse- and fine-tuning modulators. These modulators are parameters associated with the Rb-E2F network; given the same parameter perturbation degree, they induce large- and small-scale changes (coarse- and fine-tuning, respectively) in Th_E2F_.^20^ Examples of coarse-tuning (strong) modulators of deep and shallow quiescence include p21 and cyclin D, respectively^20^. However, our modeling and experimental tests were based on the cell model of rat embryonic fibroblasts (REF_s_)^4,18–21^. Do other cells under different conditions use the same or different groups of strong modulators to control quiescence depth? In this work, we sought to answer this question by modeling analysis of the behaviors of a large library of theoretical quiescent cell types and conditions. Our analysis suggests that two alternative groups of strong quiescence-depth modulators can be used across different quiescent cell types and conditions, and the choice of which modulator group to use is determined by the Rb protein level and dephosphorylation rate.

## RESULTS

### A wide and heterogeneous array of parameter conditions can seed quiescent cells

We set out to examine whether the same or case-specific strategy for quiescence-depth control is employed by different cell types under different cellular conditions. Following our previous study in REF cells^20^, we could identify the corresponding course- and fine-tuning parameters of quiescence depth by simulating the effects of parameter changes from their base values in each scenario. It is impractical, however, to obtain the actual parameter values associated with each and every cell type and condition. As such, we started by generating a large library of “seed” models, each with a random parameter set and representing a hypothetical cell type or condition.

To generate the seed model library for quiescent cells, we consider that i) a functional Rb-E2F bistable switch is required for the proper separation and transition between the two cell fates of quiescence and proliferation^4,18^, and ii) the parameter values associated with the Rb-E2F bistable switch can vary from one seed model to another within a large range. Correspondingly, we started each hypothetical model with our previously established Rb-E2F switch ODE framework^18,20^ (Table S1), with the values of the 28 model parameters randomly assigned log-uniformly within the 0.1-10x range from their original base values (Table S2). That is, we assumed that the parameter values under different cell types or conditions could vary up to 100-fold (10 times lower or higher compared to those in our previous REF cell-based model^18,20^).

Next, to identify those models that generate a functional Rb-E2F bistable switch, we simulated each model, with its randomly assigned parameter set, over 500 serum concentrations (log-uniformly distributed from 0.01 to 50) from two initial conditions (E2F-off and E2F-on, respectively; see Table S1). For those models that generated a steady-state outcome (see Methods for details), we then determined whether they generated a functional Rb-E2F bistable switch according to the three criteria established in our previous work^19^: (1) switch-like, (2) bistable, and (3) resettable (see Methods for details). In addition, we required the Th_E2F_ in the seed models to fall in a range (0.5-10 μM, criterion 4) that we considered meaningful for normal quiescent cells. In this manner, we generated 30,000 Rb-E2F models, each with a different parameter set, that fit the above 4 criteria (see Fig. S1 for the success rate change after each criterion). These 30,000 functional “seeds” models were used for subsequent analyses, each representing a theoretical condition (indicated by the corresponding seeding parameter set) at which a healthy cell may reside. As a group, there were no apparent patterns in the seeding parameter values (Fig. S2). This result suggested that quiescent cells may, at least theoretically, be associated with highly heterogeneous parameter conditions.

### Strong modulators coarse tune quiescence depth in a seeding condition-dependent manner

At each of the 30,000 seed models, we simulated the sensitivity of Th_E2F_ changes due to changes in each model parameter from its seed value. Specifically, we varied each parameter from 10-fold lower than its seed value up to 10-fold higher, while keeping other model parameters unperturbed. With each new parameter value, we first examined whether the resultant model still maintained a functional Rb-E2F bistable switch (i.e., criteria 1-3 above). If yes, we recorded the resultant Th_E2F_. As such, we examined whether and how a given parameter change may alter the Th_E2F_ of the Rb-E2F bistable switch, and thus, the quiescence depth. We performed this analysis for each of the 28 model parameters (Table S2) in a seed model and generated a corresponding “spider” graph. We then repeated this process for each of the 30,000 seed models.

Fig. 1a shows the spider graphs of 2 representative seed models. In each spider graph, we can see that parameters with a steep slope generated a larger fold change in Th_E2F_ (y-axis) than parameters with a low slope, given the same degree of parameter perturbations (fold change in parameter values; x-axis). That is, the slope steepness indicates the parameter “strength” in driving deep or shallow quiescence. Strong parameters are more sensitive than weak parameters – given the same degree of parameter perturbations, strong parameters induce larger changes in quiescence depth. To induce the same level of quiescence-depth change, a smaller degree of perturbations is needed for strong parameters than weak ones. As seen in Fig 1a, parameter strengths vary depending on the seed models. For example, the strengths of *d_CD_* and *k_RE_* were similar in one spider graph but very different in another, and the strengths of *k_M_* and *k_CDS_* even flipped between the two spider graphs.

**Figure 1.**
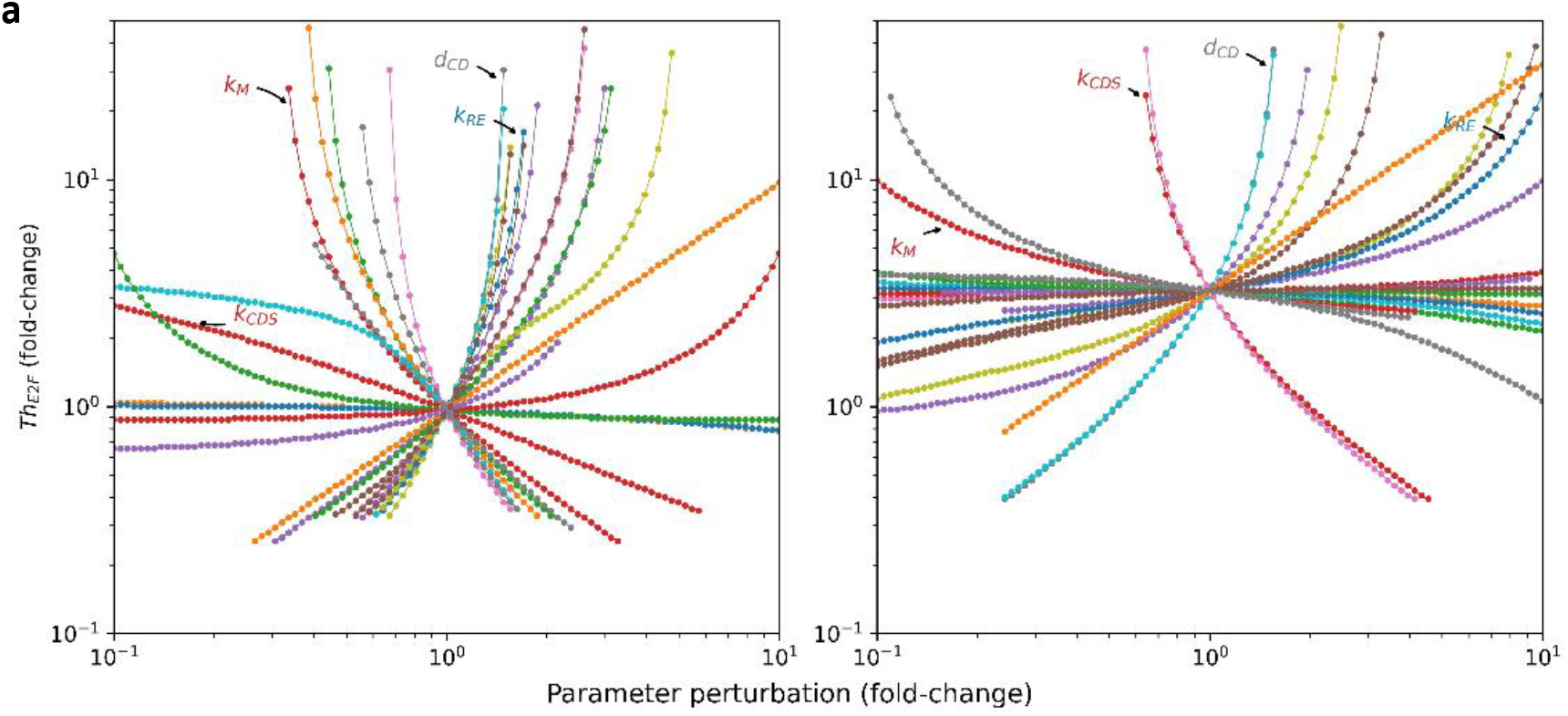
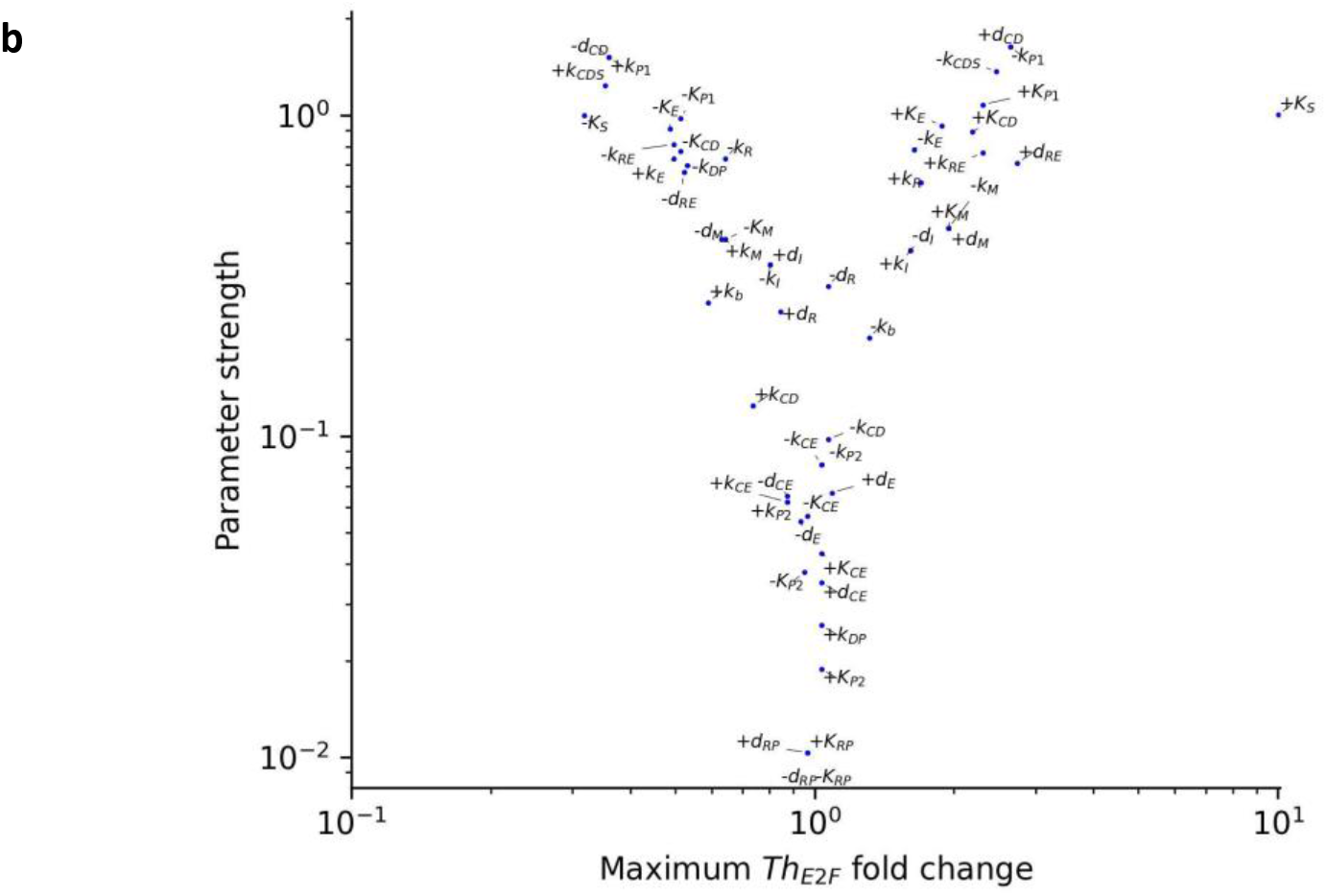
Parameter spider graph and strength. **a)** Spider graph. Each curve represents the induced fold change of Th_E2F_ (y-axis) resulting from the perturbation of the seeding parameter value (fold-change, x-axis). The curve stops where further parameter perturbations break the functional Rb-E2F bistable switch. **b)** Parameter strength vs. maximum effect. The y-axis indicates the median parameter strength across 30,000 seeding parameter sets. The x-axis indicates the median value of the induced maximum Th_E2F_ fold changes by perturbing a parameter across the 30,000 seeding parameter sets. The + and - signs before a parameter name indicate the directions (increasing and decreasing values, respectively) of the parameter perturbation (same below).

When perturbed away from its seeding value, a parameter may fail to generate a functional Rb-E2F bistable switch. This occurred when the model simulation with the new parameter value did not generate a steady-state outcome (21%) or did not meet the seeding criteria of (1) switch-like, (2) bistable, and (3) resettable (with frequencies of 19%, 51%, and 8%, respectively). In general, we found that stronger parameters tended to tolerate smaller fold changes from their seed values than weaker parameters before breaking the Rb-E2F bistable switch (Fig. S3). That said, there still appears to be a positive correlation between parameter strength and the maximum degree of induced quiescence-depth change, where strong parameters are generally able to push cells to further deeper (or shallower) quiescence in the end than weak parameters (Fig. 1b).

### Two alternative groups of strong modulators of quiescence depth

Although the parameter strengths varied from one seed model to another, those top-ranked (strongest) parameters across the 30,000 seed models appeared to converge at a small parameter group (Fig. 2a). Indeed, the membership of the top 10 strongest parameters in driving deep or shallow quiescence quickly became apparent by cumulative ranking (Fig. 2b). These top 10 strongest parameters by cumulative ranking were also consistent with those ranked by median parameter strength (as seen in the aggregated spider graph, Fig. 2c). The only exception was *Ks*, which was one of the top 10 ranked in increasing Th_E2F_ by median parameter strength but not by cumulative ranking.

**Figure 2.**
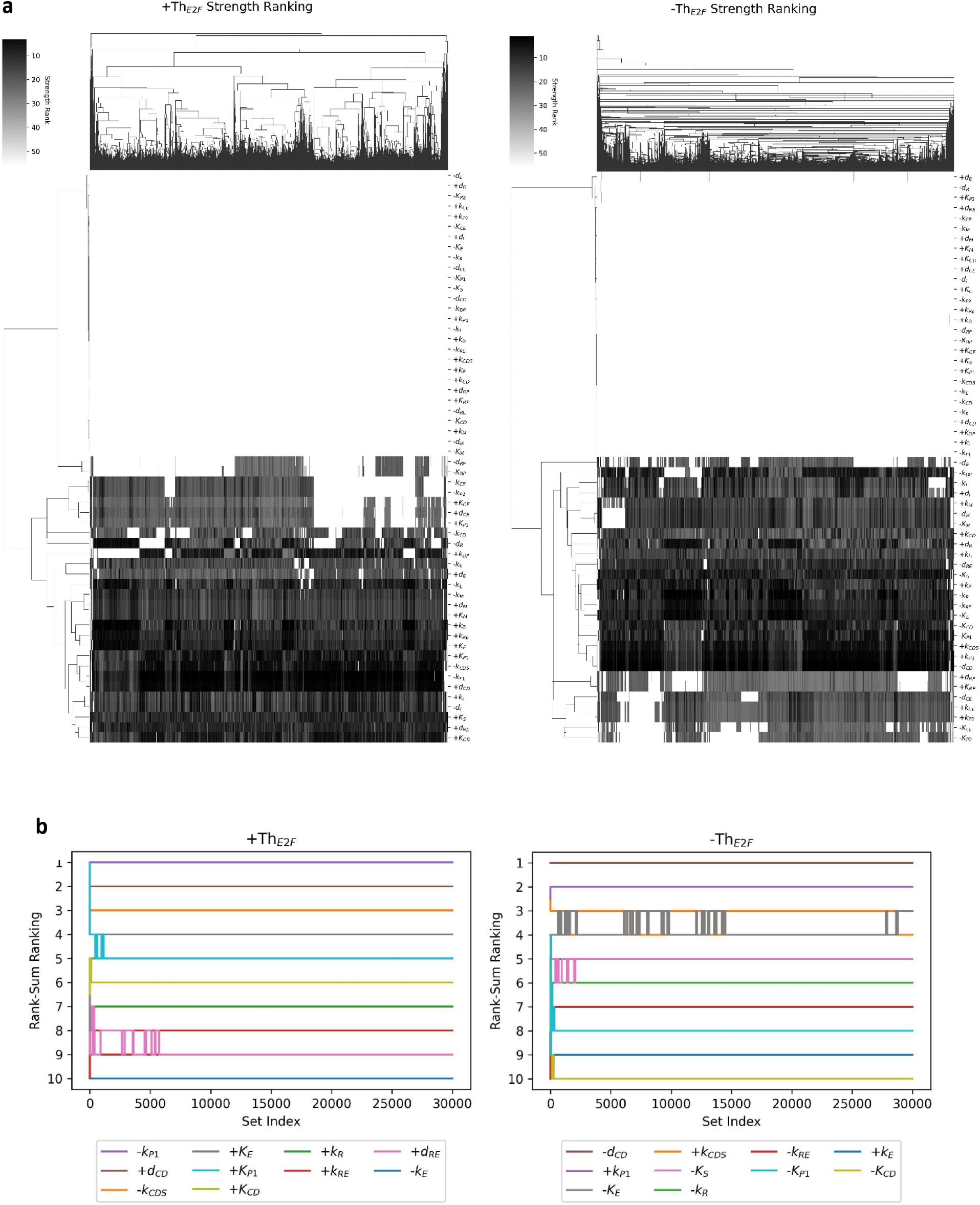
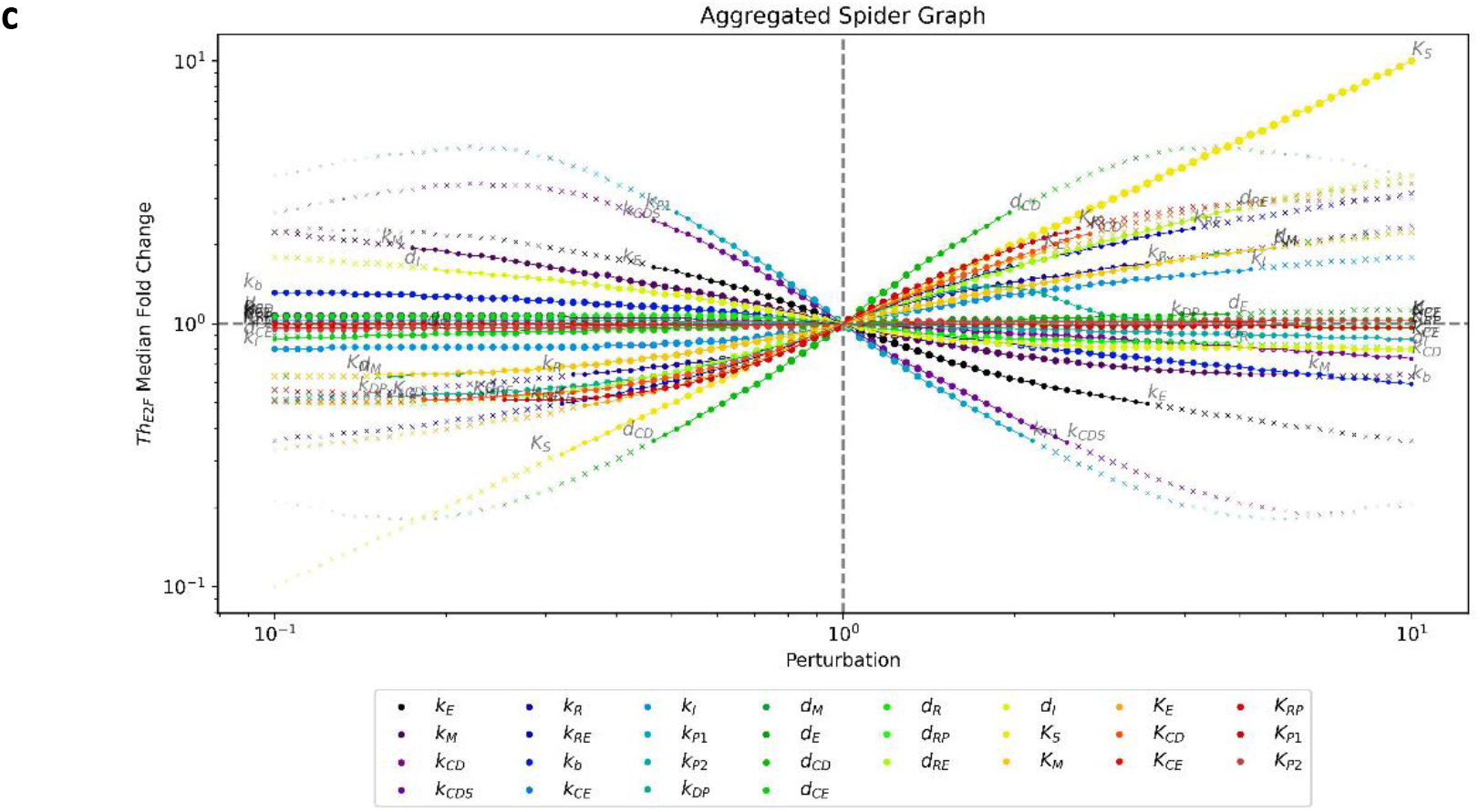
Strong modulators (parameters) of quiescence depth. **a)** Heatmap showing the parameter ranking (see the grayscale bar) across 30,000 quiescent cell seeding models (columns) with two-way hierarchical clustering. +Th_E2F_, increasing Th_E2F_; -Th_E2F_, decreasing Th_E2F_ (same below). **b)** The change in the top 10 strongest parameters in increasing (left) and decreasing (right) Th_E2F_ by cumulative ranking, along with an increasing number (x-axis) of seeding models. **c)** Aggregated spider graph. Each curve represents the median value of the Th_E2F_ fold changes across the 30,000 seeding parameter sets (y-axis) induced by the parameter perturbation (fold-change over the seeding value, x-axis). Parameters with fewer than half (n=15,000) of parameter sets surviving (without breaking the functional Rb-E2F bistable switch) at a perturbation are marked as “x”. Point sizes denote the ratio of surviving parameter sets.

We next focused on the top 10 strongest parameters by cumulative ranking and examined how they were utilized by different quiescent cells to regulate quiescence depth. We used UMAP to analyze the strength distributions of the 10 strongest parameters in increasing and decreasing Th_E2F_, respectively, across the 30,000 quiescent seed models (i.e., in two 10 × 30,000 matrices, each data point representing the strength value of a parameter in a seed model). Two distinct clusters were identified in each case (Fig 3).

**Figure 3.**
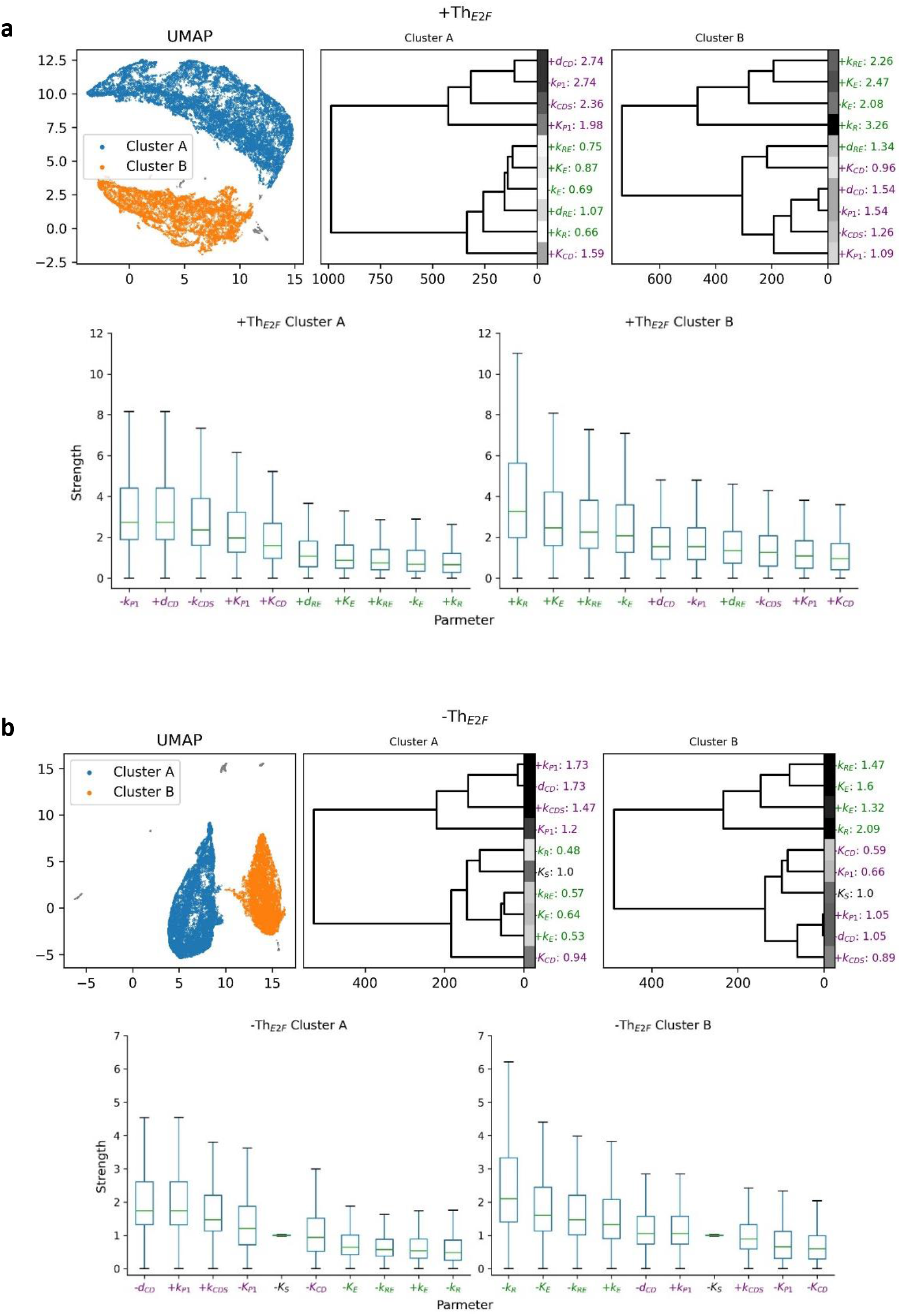
Two alternative groups of strong modulators of quiescence depth. **a)** UMAP clusters of the top 10 parameters (strength values) in increasing Th_E2F_ across 30,000 seed models (parameter sets). Cluster labels (A and B) were assigned to parameter sets using DBSCAN, and hierarchical clustering further separated parameters within each UMAP cluster according to strength values. Medians of parameter strengths across the seed models within each UMAP cluster are shown by the parameter name. Boxplots show the strength value distributions of parameters across the seed models within each UMAP cluster. **b)** UMAP clusters of the top 10 parameters (strength values) in decreasing Th_E2F_ across 30,000 seed models. See a) for details.

In the two UMAP clusters that drive deep quiescence, one cluster (cluster A, Fig 3a) exhibits a pattern in which the parameters related to the G1 cyclin/cdk activities (*d_CD_, k_CDS_, K_CD_, k_P1_*, and *K_P1_*; in purple, Fig. 3a) had higher strengths (median values 1.59-2.74) than those (median values 0.66-1.07) of parameters related to the Rb/E2F complex (*k_RE_, d_RE_, k_E_, K_E_*, and *kR*; in green, Fig. 3a). We note that this pattern is the same as what we observed previously in our studies with REF cells, in which adjusting the G1 cyclin/cdk-related parameters is the most effective in driving deep quiescence.

Interestingly, the other UMAP cluster (cluster B, Fig 3a) exhibits an opposite pattern of cluster A: the parameters related to the Rb/E2F complex (*k_RE_, d_RE_, k_E_, K_E_*, and *k_R_*; in green, Fig. 3a) exhibited higher strengths (median values 1.34-3.26) than those (median values 0.96-1.54) of parameters related to the G1 cyclin/cdk activities (*d_CD_, k_CDS_, K_CD_, k_P1_*, and *K_P1_*; in purple, Fig. 3a). That is, in the corresponding group of quiescent cells, adjusting the Rb/E2F complex-related parameters was more effective than adjusting G1 cyclin/cdk-related parameters in driving deep quiescence.

Similarly, in the two UMAP clusters that drive shallow quiescence, one group of quiescent cells (cluster A, Fig 3b) preferentially used G1 cyclin/cdk-related parameters as the strong modulators (median strength values 0.94-1.73; in purple, Fig. 3b) over Rb/E2F-related parameters (median strength values 0.48-0.64; in green, Fig. 3b). In contrast, another group of quiescent cells (cluster B, Fig 3b) preferred to use Rb/E2F-related parameters (median strength values 1.32-2.09; in green, Fig 3b) as the more effective ones than G1 cyclin/cdk-related parameters (median strength values 0.59-1.05; in purple, Fig 3b) in driving shallow quiescence.

### The Rb level and dephosphorylation rate determine which strong modulator group to use to regulate quiescence depth

Given that there are two alternative strong modulator groups (G1 cyclin/cdk-related and Rb/E2F complex-related) to alter quiescence depth, we wondered what cellular features determine which strong modulator group is employed. To this end, we constructed a decision tree model to identify the features (parameters) in the Rb-E2F bistable switch that classify one UMAP cluster from the other. Ultimately, we identified *k_R_, k_DP_*, and *d_R_* (the rates of Rb synthesis, dephosphorylation, and degradation, respectively) to be the three critical features.

Specifically, the 30,000 theoretical quiescent cells with a functional Rb-E2F bistable switch can be primarily classified into two groups (A and B, Fig. 4) with high accuracy (ROC plots, Fig. 4). One group of quiescent cells (Group A) is enriched with large *k_R_* but small *k_DP_* and *d_R_* values (Fig. 4), and thus features a large amount of Rb with a low dephosphorylation rate. These cells favor G1 cyclin/cdk-related parameters as strong modulators in regulating quiescence depth (Cluster A, Fig. 3). In driving deep quiescence by increasing Th_E2F_ (Fig. 4a), for example, 56% of the 30,000 quiescent cell seeds are classified as Group A with 84% accuracy based on their relatively large *k_R_* values (>= 0.14) alone. We note that our original model related to REF cells belongs to this group (*k_R_* = 0.18; Table S2). For those cells with *k_R_* < 0.14 to still favor G1 cyclin/cdk-related modulators, their best chance is to have small *k_DP_* and *d_R_* values (<2.6 and <0.056, respectively, Fig. 4a). This latter case explains 11% of the 30,000 quiescent cell seeds with 81% accuracy. A similar pattern holds in driving shallow quiescence by decreasing Th_E2F_. As seen in the tree in Fig. 4b, quiescent cells with a large *k_R_* (>= 0.15) and small *dR* (< 0.092), for example, are classified into Group A (favoring G1 cyclin/cdk-related parameters to drive shallow quiescence) with 91% accuracy, which accounts for 34% of the 30,000 quiescent cell seeds.

**Figure 4.**
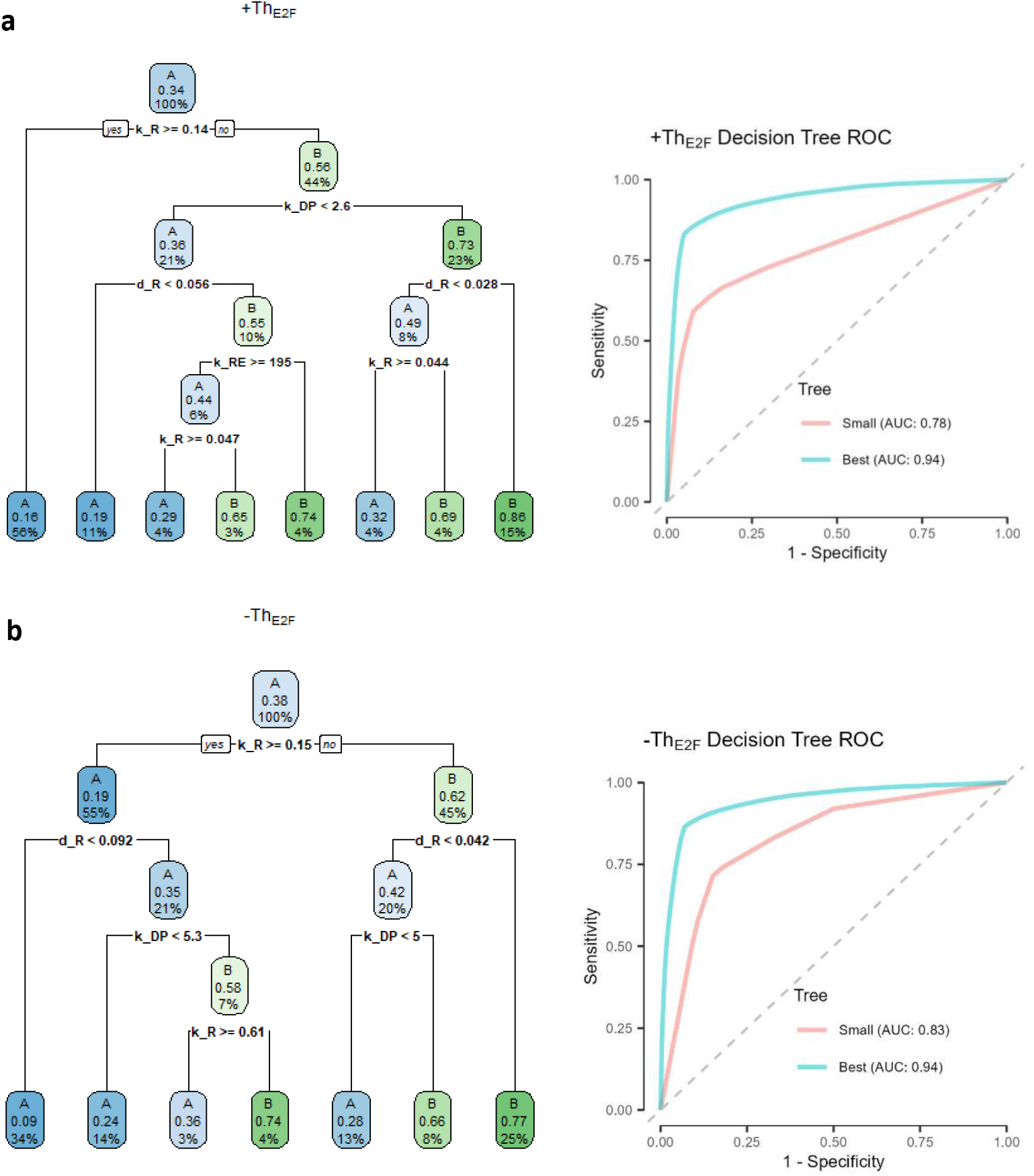
Decision tree analysis of the two alternative strong modulator groups in **a**) increasing Th_E2F_ and **b**) decreasing Th_E2F_. The small tree (left) shows the top splits (≤ 5 layers) of the best model (with the least cross-validation error). The AUC/ROC of the small and best tree models are shown to the right.

In contrast, the other group in the 30,000 quiescent cells is enriched with small *k_R_* but large *d_R_* and *k_DP_* values (Group B, Fig 4), thus featuring a small amount of Rb with a high dephosphorylation rate. These cells favor Rb/E2F complex-related parameters, instead of G1 cyclin/cdk-related, as strong modulators in regulating quiescence depth (Fig 4). In driving deep quiescence by increasing Th_E2F_ (Fig. 4a), for example, 44% of the 30,000 quiescent cell seeds are classified as Group B (favoring Rb/E2F complex-related modulators) with 56% accuracy based on their relatively small *k_R_* values (< 0.14) alone. The accuracy of Group B classification increased to 73% with the additional condition of *k_DP_* > 2.6, and further increased to 86% with *d_R_* > 0.028 (Fig. 4a). A similar pattern holds in driving shallow quiescence by decreasing Th_E2F_. As seen in the tree in Fig. 4b, quiescent cells with a small *k_R_* (< 0.15) and large *d_R_* (> 0.042), for example, are classified as Group B (favoring Rb/E2F complex-related parameters to drive shallow quiescence) with 77% accuracy, which accounts for 25% of the 30,000 quiescent cell seeds.

To test the model prediction above, we generated 5,000 additional quiescent seed models (following the same procedure used to generate the previous 30,000 seed model training set). We then applied the decision tree algorithm (Fig. 4) to this new test set. The accuracy of the tree model prediction was 80.8% and 85.6% for increasing Th_E2F_ and driving deep quiescence with the small and best tree models, respectively, and the accuracy of the tree model prediction was 80.1% and 85.4% for decreasing Th_E2F_ and driving shallow quiescence with the small and best tree models, respectively, in the new test set.

## DISCUSSION

The regulation of quiescence depth is critical to the proper balance and transition between cellular quiescence and proliferation. Understanding the corresponding regulatory mechanisms is needed for the development of effective treatment strategies against hypo- or hyper-proliferation diseases such as aging and cancer. Based on earlier work including ours^4,18–20,22^, we propose that a functional Rb-E2F bistable switch controls the quiescence-to-proliferation transition under normal physiological conditions across different cell types and that the activation threshold of this bistable switch (Th_E2F_) regulates quiescence depth.

By coupling modeling and experimental tests, we previously identified groups of coarse- and fine-tuning modulators of Th_E2F_ and thus quiescence depth in REF cells^20^. However, as revealed in this study, a wide and heterogeneous array of parameter conditions can support a functional Rb-E2F bistable switch, and each may represent a theoretical quiescence cell under a given environmental condition. Do these cells employ the same or distinct groups of quiescence-depth modulators?

Our further analysis suggested that among these vastly heterogeneous quiescent cells (with no apparent parameter patterns across the corresponding seed models, Fig. S2), two alternative groups of strong quiescence-depth modulators emerge. One group is related to the G1 cyclin/cdk activities (*d_CD_, k_CDS_, K_CD_, k_P1_, K_P1_*; in purple, Fig. 3), and another group is related to the Rb/E2F complex (*k_RE_, d_RE_, k_E_, K_E_, k_R_*, (*d_RE_);* in green, Fig. 3). Different quiescent cells (among the 30,000 seed models) appear to use one group or the other as strong modulators to regulate quiescence depth.

By building a machine learning model (decision tree), we were able to identify which features associated with the Rb-E2F bistable switch determine which of the two alternative groups of strong quiescence-depth modulators a cell will utilize. Quiescent cells with large *k_R_* but small *k_DP_* and *d_R_* values (Group A, Fig. 4) favor G1 cyclin/cdk-related parameters as strong modulators in regulating quiescence depth, whereas quiescent cells with small *k_R_* but large *k_DP_* and *d_R_* values (Group B, Fig. 4) favor Rb/E2F complex-related strong modulators. Retrospectively, this result makes some intuitive sense. In Group A cells, large *k_R_* and small *d_R_* lead to a high Rb level, which tends to desensitize Rb/E2F-related parameter changes; they have small *k_DP_* (small dephosphorylation baseline) and are thus likely sensitive to G1 cyclin/cdk (phosphorylation)-related modulators. In Group B cells, small *k_R_* and large *d_R_* lead to a low Rb baseline and thus they become sensitive to Rb/E2F-related parameter changes; they have large *k_DP_* (high dephosphorylation level), which may “buffer” the effects caused by G1 cyclin/cdk (phosphorylation)-related modulators. We note that the REF cell model used in our previous studies belongs to Group A. What other cell types belong to Group B, and whether we can convert cells from Group A to Group B and thus change their quiescence-depth modulator preference by following the decision rules (Fig. 4), are examples of interesting questions generated by this current modeling analysis to be tested in future experiments.

## Methods

### Model simulation

Ordinary differential equations (ODEs) of the Rb-E2F bistable switch model used for time-course simulation were described in our previous work^18,20^ (Table S1). 100 time points over the interval [0-1000] were simulated using SciPy’s odeint (mxstep=100000, rtol=1e^−6^, and atol=1e^−12^). E2F-OFF initial conditions were determined as the steady-state values simulated from the default initial condition (initialized at 0 for all concentrations except [R]=0.55 *μ*M, and [I]=0.5 *μ*M), using model-specific parameter sets. E2F-ON initial conditions were determined as the steady-state values simulated from the E2F-OFF initial condition using the corresponding model-specific parameter set. Concentrations and parameters were converted to counts before simulation using the equation *Counts* = 6.02*E*23 * *V_Cell_* * [*X*] where the assumed cell volume, V_cell_, is 1e^−12^ L and [X]=1E6 is the concentration unit. A steady state was determined to be reached when the difference between the E2F counts at the last two time points was ≤ 50 as simulated from E2F-OFF and -ON initial conditions, respectively. Time course simulations were run for each of 500 serum concentrations from [S] = 0.01 to 50 *μ*M, log-uniformly spaced. Simulation and data processing were performed using high-performance computing (HPC) at the University of Arizona.

### Seed model library (parameter sets) generation

Random parameter sets were generated by multiplying each parameter in the base parameter set (Table S2) by a value sampled from 0.1 to 10, distributed log-uniformly. Serum stimulations were simulated on each new parameter set. Parameter sets that pass all criteria 1-4 below were considered valid seeding parameter sets. Bistable regions were detected when the difference between the steady-state values of E2F simulated from E2F-OFF and E2F-ON initial conditions exceeded a tolerance of 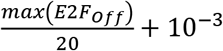 (with max(*E2F_off_*) being the maximum E2F level simulated from the E2F-OFF initial condition). Parameter sets were determined to be unstable and discarded if more than 5 of the simulated serum concentrations did not reach E2F steady state, and otherwise determined to generate a steady-state outcome.

1. **Switch-like**: The difference between the minimum and maximum E2F levels simulated from the E2F-OFF initial condition is > 0.1 *μM*.
2. **Bistable**: The width of the bistable region is ≥ 0.2 *μM*.
3. **Resettable**: The difference between E2F steady states simulated from E2F-OFF and E2F-ON initial conditions approaches 0 as [S] approaches 0
4. **Biologically sound threshold**: Th_E2F_ ∈ [0.5 *μM*, 10 *μM*]

### Parameter perturbation and parameter strength calculation

Each parameter value of a seeding parameter set was individually perturbed by multiplying by 100 values from 0.1 to 10, distributed log-uniformly. Th_E2F_ values were standardized using their fold change of Th_seed_ (Th_E2F_ at the seeding parameter set), such that 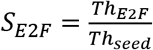. Parameter strength was calculated as 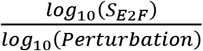 at the point where log_10_(S_E2F_) was one half of the maximum log_10_(S_E2F_).

### UMAP and hierarchical clustering

The strength values of the top 10 strongest parameters were dimensionally reduced using uniform manifold approximation and projection (UMAP). Clusters were assigned labels using Scikit-learn’s implementation of the density-based spatial clustering of applications with noise (DBSCAN) algorithm. Dendrograms were calculated from Scikit-learn’s implementation of hierarchical clustering using Ward’s method and the Euclidean metric. Hierarchically clustered heatmaps were created using the Seaborn package.

### Decision Tree Analysis

Decision trees were trained on 30,000 seeding parameter sets using the R package rpart. Cluster assignment was used as labels, and parameter values as predictors. The remaining 5,000 parameter sets were used as a hold-out for validation. The “best” models were selected by minimizing the 10-fold cross-validation error and “small” interpretable models were selected to show the top splits (≤ 5 layers). ROC curves were calculated using the ROCR R package.

## REFERENCE

1. Cheung, T.H. & Rando, T.A. Molecular regulation of stem cell quiescence. Nat Rev Mol Cell Biol 14, 329–40 (2013).

2. Legesse-Miller, A. et al. Quiescent fibroblasts are protected from proteasome inhibition-mediated toxicity. Mol Biol Cell 23, 3566–81 (2012).

3. Coller, H.A., Sang, L. & Roberts, J.M. A New Description of Cellular Quiescence. PLoS Biol 4, e83 (2006).

4. Yao, G. Modelling mammalian cellular quiescence. Interface Focus 4, 20130074 (2014).

5. Augenlicht, L.H. & Baserga, R. Changes in the G0 state of WI-38 fibroblasts at different times after confluence. Exp Cell Res 89, 255–62 (1974).

6. Martin, R.G. & Stein, S. Resting state in normal and simian virus 40 transformed Chinese hamster lung cells. Proc Natl Acad Sci U S A 73, 1655–9 (1976).

7. Owen, T.A., Soprano, D.R. & Soprano, K.J. Analysis of the growth factor requirements for stimulation of WI-38 cells after extended periods of density-dependent growth arrest. J Cell Physiol 139, 424–31 (1989).

8. Miska, D. & Bosmann, H.B. Existence of an upper-limit to elongation of the prereplicative period in confluent cultures of C3H/10T 1/2 cells. Biochem Biophys Res Commun 93, 1140–5 (1980).

9. Yanez, I. & O’Farrell, M. Variation in the length of the lag phase following serum restimulation of mouse 3T3 cells. Cell Biol Int Rep 13, 453–62 (1989).

10. Gunther, G.R., Wang, J.L. & Edelman, G.M. The kinetics of cellular commitment during stimulation of lymphocytes by lectins. J Cell Biol 62, 366–77 (1974).

11. Burmer, G.C., Rabinovitch, P.S. & Norwood, T.H. Evidence for differences in the mechanism of cell cycle arrest between senescent and serum-deprived human fibroblasts: heterokaryon and metabolic inhibitor studies. Journal of cellular physiology 118, 97–103 (1984).

12. Bucher, N.L. Regeneration of Mammalian Liver. Int Rev Cytol 15, 245–300 (1963).

13. Roth, G.S. & Adelman, R.C. Age-dependent regulation of mammalian DNA synthesis and cell division in vivo by glucocorticoids. Exp Gerontol 9, 27–31 (1974).

14. Rodgers, J.T. et al. mTORC1 controls the adaptive transition of quiescent stem cells from G0 to G(Alert). Nature 510, 393–6 (2014).

15. Llorens-Bobadilla, E. et al. Single-Cell Transcriptomics Reveals a Population of Dormant Neural Stem Cells that Become Activated upon Brain Injury. Cell Stem Cell 17, 329–40 (2015).

16. Rodgers, J.T., Schroeder, M.D., Ma, C. & Rando, T.A. HGFA Is an Injury-Regulated Systemic Factor that Induces the Transition of Stem Cells into GAlert. Cell Rep 19, 479–486 (2017).

17. Lee, G. et al. Fully reduced HMGB1 accelerates the regeneration of multiple tissues by transitioning stem cells to GAlert. Proc Natl Acad Sci U S A 115, E4463–E4472 (2018).

18. Yao, G., Lee, T.J., Mori, S., Nevins, J.R. & You, L. A bistable Rb-E2F switch underlies the restriction point. Nat Cell Biol 10, 476–82 (2008).

19. Yao, G., Tan, C., West, M., Nevins, J.R. & You, L. Origin of bistability underlying mammalian cell cycle entry. Mol Syst Biol 7, 485 (2011).

20. Kwon, J.S. et al. Controlling Depth of Cellular Quiescence by an Rb-E2F Network Switch. Cell Rep 20, 3223–3235 (2017).

21. Fujimaki, K. et al. Graded regulation of cellular quiescence depth between proliferation and senescence by a lysosomal dimmer switch. Proc Natl Acad Sci U S A 116, 22624–22634 (2019).

22. Wang, X. et al. Exit from quiescence displays a memory of cell growth and division. Nature communications 8, 321 (2017).

