## Supplemental Figures and Tables for "Identifying Strong Modulators of Cellular Quiescence Depth Across Different Quiescent Cells and Conditions"

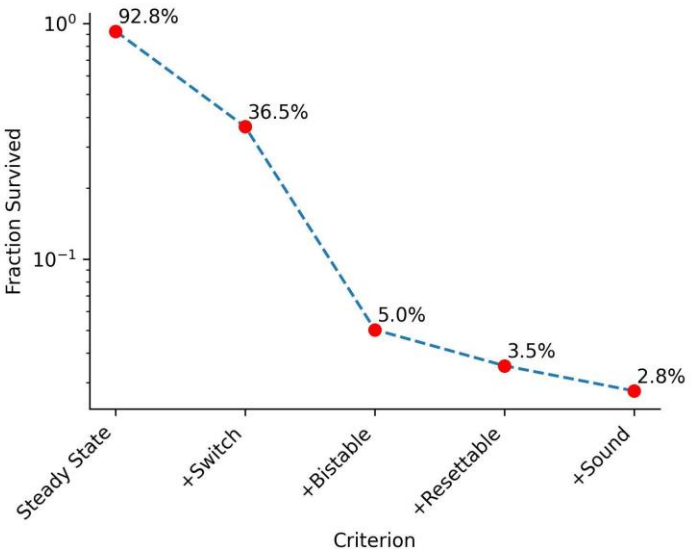


**Figure S1.** The survival rate of random parameter sets after each seeding criterion. Shown is the sequential success rate (y-axis, in log scale) of the starting random parameter sets that passed each criterion for a functional Rb-E2F bistable switch "seed" (x-axis) to be considered as a theoretical quiescent cell. See Methods for the description of each criterion in detail.

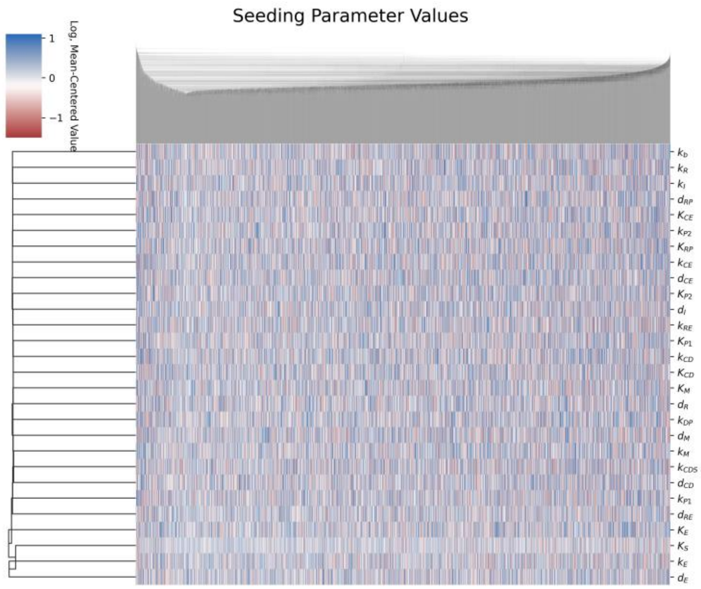


**Figure S2.** Heterogeneity in seeding parameter sets for theoretical quiescent cells. The heatmap shows parameter values ($\log_{10}$ transformed and mean-centered by each row) with two-way hierarchical clustering. Each column corresponds to one of the 30,000 quiescent cell seeding models.

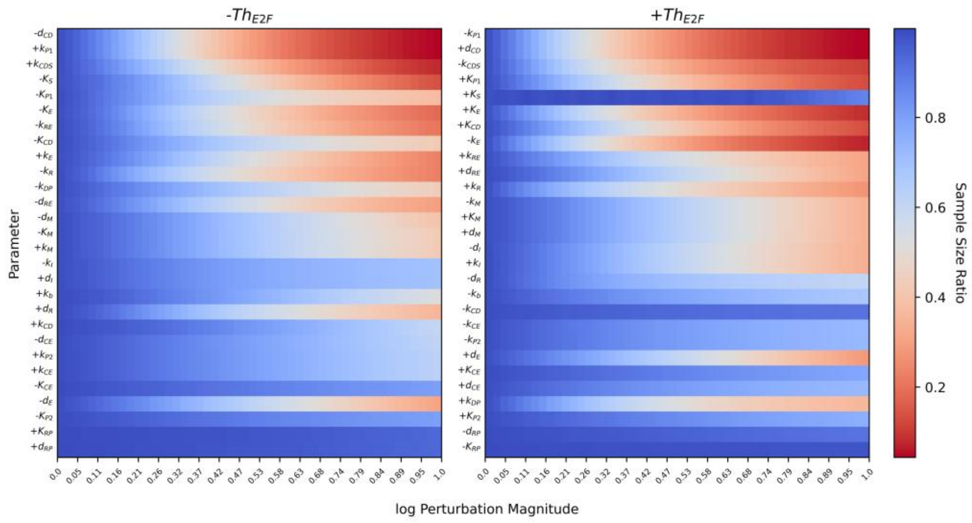


**Figure S1**. Parameter-perturbation tolerance (flexibility). Indicated by the color scheme (right) is the ratio (out of 30,000) of parameter sets surviving the seeding criteria 1-3 after perturbing a parameter away from seeding parameter values with the indicated perturbation scale (in log scale, x-axis). Parameters (rows) are ranked from top to bottom by the median strength value of each parameter across the 30,000 seeding parameter sets.

**Table S1. The Rb-E2F switch model** (adapted from Ref[20]).

| |
|---|
| $\frac{d[M]}{dt} = \frac{k_M[S]}{K_S + [S]} - d_M[M]$ |
| $\frac{d[E]}{dt} = k_E\left(\frac{[M]}{K_M + [M]}\right)\left(\frac{[E]}{K_E + [E]}\right) + \frac{k_b[M]}{K_M + [M]} + \left(\frac{k_{P1}}{K_{P1} + [I]}\right)\left(\frac{[CD][RE]}{K_{CD} + [RE]}\right) + \left(\frac{k_{P2}}{K_{P2} + [I]}\right)\left(\frac{[CE][RE]}{K_{CE} + [RE]}\right) - d_E[E] - k_{RE}[R][E]$ |
| $\frac{d[CD]}{dt} = \frac{k_{CD}[M]}{K_M + [M]} + \frac{k_{CDS}[S]}{K_S + [S]} - d_{CD}[CD]$ |
| $\frac{d[CE]}{dt} = \frac{k_{CE}[E]}{K_E + [E]} - d_{CE}[CE]$ |
| $\frac{d[R]}{dt} = k_R + \frac{k_{DP}[RP]}{K_{RP} + [RP]} - k_{RE}[R][E] - \left(\frac{k_{P1}}{K_{P1} + [I]}\right)\frac{[CD][R]}{K_{CD} + [R]} - \left(\frac{k_{P2}}{K_{P2} + [I]}\right)\frac{[CE][R]}{K_{CE} + [R]} - d_R[R]$ |
| $\frac{d[RP]}{dt} = \left(\frac{k_{P1}}{K_{P1} + [I]}\right)\frac{[CD][R]}{K_{CD} + [R]} + \left(\frac{k_{P2}}{K_{P2} + [I]}\right)\frac{[CE][R]}{K_{CE} + [R]} + \left(\frac{k_{P1}}{K_{P1} + [I]}\right)\frac{[CD][RE]}{K_{CD} + [RE]} + \left(\frac{k_{P2}}{K_{P2} + [I]}\right)\frac{[CE][RE]}{K_{CE} + [RE]} - \frac{k_{DP}[RP]}{K_{RP} + [RP]} - d_{RP}[RP]$ |
| $\frac{d[RE]}{dt} = k_{RE}[R][E] - \left(\frac{k_{P1}}{K_{P1} + [I]}\right)\frac{[CD][RE]}{K_{CD} + [RE]} - \left(\frac{k_{P2}}{K_{P2} + [I]}\right)\frac{[CE][RE]}{K_{CE} + [RE]} - d_{RE}[RE]$ |
| $\frac{d[I]}{dt} = k_I - d_I[I]$ |

*Model variables*:
S: serum concentration
M: Myc
E: E2F
CD: Cyclin D/Cdk4,6
CE: Cyclin E/Cdk2
R: Rb family proteins
RP: Phosphorylated Rb
RE: Rb-E2F complex
I: Cdk inhibitors

**Table S2. Model parameters** (adapted from (adapted from Ref[20]).

| Symbol | Values | Description |
|---|---|---|
| $k_M$ | 1.0 µM/hr | Rate constant of Myc synthesis driven by growth factors |
| $k_E$ | 0.4 µM/hr | Rate constant of E2F synthesis driven by Myc and E2F |
| $k_b$ | 0.003 µM/hr | Rate constant of E2F synthesis driven by Myc alone |
| $k_{CD}$ | 0.03 µM/hr | Rate constant of CycD synthesis driven by Myc |
| $k_{CDS}$ | 0.45 µM/hr | Rate constant of CycD synthesis driven by growth factors |
| $k_{CE}$ | 0.35 µM/hr | Rate constant of CycE synthesis driven by E2F |
| $k_R$ | 0.18 µM/hr | Rate constant of Rb constitutive synthesis |
| $k_I$ | 0.15 µM/hr | Rate constant of Cdk inhibitor synthesis |
| $k_{DP}$ | 3.6 µM/hr | Dephosphorylation rate constant of Rb by phosphatases |
| $k_{RE}$ | 180 µM/hr | Association rate constant of Rb and E2F |
| $K_S$ | 0.5 µM | Michaelis-Menten parameter for CycD and Myc synthesis by growth factors |
| $K_E$ | 0.15 µM | Michaelis-Menten parameter for CycE and E2F synthesis by E2F |
| $K_M$ | 0.15 µM | Michaelis-Menten parameter for CycD and E2F synthesis by Myc |
| $K_{RP}$ | 0.01 µM | Michaelis-Menten parameter for Rb dephosphorylation |
| $K_{CD}$ | 0.92 µM | Michaelis-Menten parameter for Rb phosphorylation by CycD/Cdk4,6 |
| $K_{CE}$ | 0.92 µM | Michaelis-Menten parameter for Rb phosphorylation by CycE/Cdk2 |
| $d_M$ | 0.7/hr | Degradation rate constant of Myc |
| $d_E$ | 0.25/hr | Degradation rate constant of E2F |
| $d_{CD}$ | 1.5/hr | Degradation rate constant of CycD |
| $d_{CE}$ | 1.5/hr | Degradation rate constant of CycE |
| $d_R$ | 0.06/hr | Degradation rate constant of Rb |
| $d_{RP}$ | 0.06/hr | Degradation rate constant of phosphorylated Rb |
| $d_{RE}$ | 0.03/hr | Degradation rate constant of Rb-E2F complex |
| $d_I$ | 0.3/hr | Degradation rate constant of Cdk inhibitor p21 |
| $k_{P1}$ | 45/hr | Phosphorylation rate constant of CycD/Cdk4,6 |
| $k_{P2}$ | 45/hr | Phosphorylation rate constant of CycE/Cdk2 (Cdks) |
| $K_{P1}$ | 2 µM | Michaelis-Menten parameter for CycD/Cdk4,6 activities affected by Cdk inhibitors |
| $K_{P2}$ | 2 µM | Michaelis-Menten parameter for CycE/Cdk2 activities affected by Cdk inhibitors |

**Table S3. Parameter cumulative ranking**

| $+Th_{E2F}$ | Rank | $-Th_{E2F}$ | Rank |
|---|---|---|---|
| $-d_{CD}$ | 1 | $-k_{P1}$ | 1 |
| $+k_{P1}$ | 2 | $+d_{CD}$ | 2 |
| $-K_E$ | 3 | $-k_{CDS}$ | 3 |
| $+k_{CDS}$ | 4 | $+K_E$ | 4 |
| $-K_S$ | 5 | $+K_{P1}$ | 5 |
| $-k_R$ | 6 | $+K_{CD}$ | 6 |
| $-k_{RE}$ | 7 | $+k_R$ | 7 |
| $-K_{P1}$ | 8 | $+k_{RE}$ | 8 |
| $+k_E$ | 9 | $+d_{RE}$ | 9 |
| $-K_{CD}$ | 10 | $-k_E$ | 10 |
| $-d_{RE}$ | 11 | $+K_S$ | 11 |
| $-k_{DP}$ | 12 | $-d_I$ | 12 |
| $+d_R$ | 13 | $+k_{DP}$ | 13 |
| $-d_M$ | 14 | $-k_M$ | 14 |
| $-K_M$ | 15 | $+k_I$ | 15 |
| $-k_I$ | 16 | $+d_M$ | 16 |
| $+k_b$ | 17 | $+K_M$ | 17 |
| $+k_M$ | 18 | $-d_R$ | 18 |
| $+d_I$ | 19 | $-k_b$ | 19 |
| $+k_{CD}$ | 20 | $+d_E$ | 20 |
| $-d_{CE}$ | 21 | $-k_{CD}$ | 21 |
| $+k_{CE}$ | 22 | $-k_{P2}$ | 22 |
| $+k_{P2}$ | 23 | $-k_{CE}$ | 23 |
| $-d_E$ | 24 | $+d_{CE}$ | 24 |
| $-K_{P2}$ | 25 | $+K_{P2}$ | 25 |
| $-K_{CE}$ | 26 | $+K_{CE}$ | 26 |
| $+K_{RP}$ | 27 | $-K_{RP}$ | 27 |
| $+d_{RP}$ | 28 | $-d_{RP}$ | 28 |
